# Gene expression is associated with virulence in murine macrophages infected with *Leptospira* spp

**DOI:** 10.1101/619023

**Authors:** Erivelto Corrêa de Araújo Junior, Leandro Encarnação Garcia, Matheus Janeck Araújo, Itamar Souza Oliveira-Junior, Daniel Robert Arnold, Flavia Lombardi Lopes, Márcia Marinho

**Affiliations:** Department of Production and Animal Health, São Paulo State University (Unesp), School of Veterinary Medicine, Araçatuba, SP, Brazil; Department of Surgery, Discipline of Anesthesia, Pain and Intensive Medicine, Federal University of São Paulo, São Paulo, Brazil

**Keywords:** Leptospirosis, macrophages, immune response, microarray

## Abstract

*Leptospira* genus contains species that affect human health with varying degrees of pathogenicity. In this context, we aimed to evaluate the differences in modulation of host gene expression by strains of *Leptospira* with varied virulence degrees. Our data showed a high number of differentilly expressed transcripts in murine macrophages following 6h of infection with both virulent and culture-attenuated *L. interrogans* and to a lesser degree, with the saprophyte strain *L. biflexa*. That suggests that certain genes are modulated by *Leptospira* infection independent of their degree of virulence, whether others are virulence and species associated. Pathway analysis indicated that Apoptosis, ATM Signaling and Cell Cycle: G2/M DNA Damage Checkpoint Regulation were exclusively regulated following infection with the virulent strain. Results demonstrated that species and virulence play a role during host response to *Leptosppira* spp in murine macrophages.

**Author summary:** Leptospirosis is an infectious disease that is transmitted from animals to humans. It is a re-emerging neglected zoonosis that is found in a range of environments worldwide, most notably tropical regions prone to flooding. This bacteria is found in soil and water and are eliminated in the urine by rats, their natural host reservoir. Through skin contacts with the bacteria people or animals can get infected however the infection process is still poorly understood, such as the fact that different strains can cause different severity of illness. In this study, we aimed to evaluate the differences in modulation of host gene expression by strains of *Leptospira* varying in virulence. After transcriptomic analysis, the results showed a high number of differentially expressed genes after 6h of infection by virulent and attenuated *L. interrogans*, and to a lesser extent with *L. biflexa* saprophytic lineage. This suggests that RNAs are modulated after infection by *Leptospira* in macrophages, in a species and virulence related manner. It is hoped that the data produced will contribute to further our understanding on the pathogenesis of leptospirosis.

## Introduction

Leptospirosis is a zoonotic bacterial disease that occurs in different epidemiological conditions [1]. The genus *Leptospira* encompasses pathogenic and saprophytic species that differ in their ability to survive and colonize different environments and hosts [2]. *Leptospira* species are classified into three groups according to their pathogenic potential: virulent pathogenic, intermediate, and saprophytes [3]. Leptospirosis occurs mainly in vulnerable populations, including urban and rural dwellers [4] of tropical and subtropical developing countries [5–7]. It is a major public health problem, with a recent estimate of 1 million cases per year, and a mortality rate of 5 to 10% [4,8–9].

Leptospires are capable of infecting humans and many domestic and wild animals, survive and thrive in host tissues, escaping from the host’s natural defense mechanisms. Transmission is based on direct or indirect contact with the urine of carriers (mainly rodents); the disease varied from sub-clinical to most serious cases, progressing to renal failure and pulmonary hemorrhage [10–1].

Host-specific immune response against pathogenic leptospires are poorly understood, particularly regarding susceptibility resistance to infection. For decades, adaptive humoral immunity was considered as the sole player in leptospirosis, but in recent years some progress has been achieved in the fields of innate and adaptive immunity [11–14].

Murray and coworkers [15] identified a number of virulence factors, including the presence of lipopolysaccharides (LPS), heme oxygenase [16], Loa22 lipoprotein [17] and other proteins related to macrophage interaction with *Leptospira*.

Phagocytosis is one of the main mechanisms to eliminate invading microbial pathogens at the early stages of infection in individuals without acquired immunity against the infecting agent, but pathogenic *Leptospira* can escape from complement attack and phagocytosis after infection [18–20]. Pathogenic *Leptospira* is also able to survive and replicate in human macrophages, but it is killed in murine macrophages [21]. LPSs of pathogenic *Leptospira* activate human macrophages only through the Toll-like receptor 2 (TLR2) while murine macrophages are activated through TLR2 and TLR4 [13–22]. Vernon Pauillac and Merian [23] have shown that mononuclear macrophages of peripheral blood of hamster infected with a virulent variant of *Leptospira interrogans* secrete proinflammatory cytokines (TNF-α) with a Th1 (IL-12) profile in the first hour, predominating until the fourth day after infection, whereas a Th2 profile appears after 24 hours of infection. In the early course of infection, leptospires have to survive and spread in the bloodstream before causing damage to target organs [25].

In this study, we applied microarray technology to comparatively analyze early change in murine macrophages genes expression in response to *Leptospira* spp. with varied virulence, and to identify signaling pathways that play a role in an *in vitro* model of macrophageal infection.

## Results

### Data deposition

Microarray raw data files are available in Gene Expression Omnibus (GEO) and are accessible through GEO series number GSE105141 [26].

### Gene expression profile via microarray analysis

Our data analysis found 892 genes in cells infected with saprophyte, attenuated and virulent leptospirosis compared to control. According to Fig 1, pathogenic leptospires modulates 892 genes (422 up and 470 down-regulated), attenuated leptospires modulates 848 genes (400 upregulated and 448 downregulated) and saprophyte 299 genes (128 upregulated and 171 downregulated) in a filter criterion of fold change ±2 and false discovery rate (FDR)<0.05 (Fig. 1). Through treatment comparison by Venn diagram, we identify common and specific genes (Fig. 2). A total of 274 genes were common to all infected cells, despite of strains, when compared to control. Virulent and culture-attenuated infected groups groups shared 512 genes in common, while eight genes were shared between virulent and saprophyte groups and only one gene between attenuated and saprophyte infected cells (S1. Table). Average singals (log2) of samples were hierarchically clustered using Pearson correlation and complete-linkage and it was observed again a clustering of samples based on species and virulence, with the virulent and culture-attenuated strains clustering closer together, followed bt the saprophyte strain (Fig. 3).

**Figure 1.**
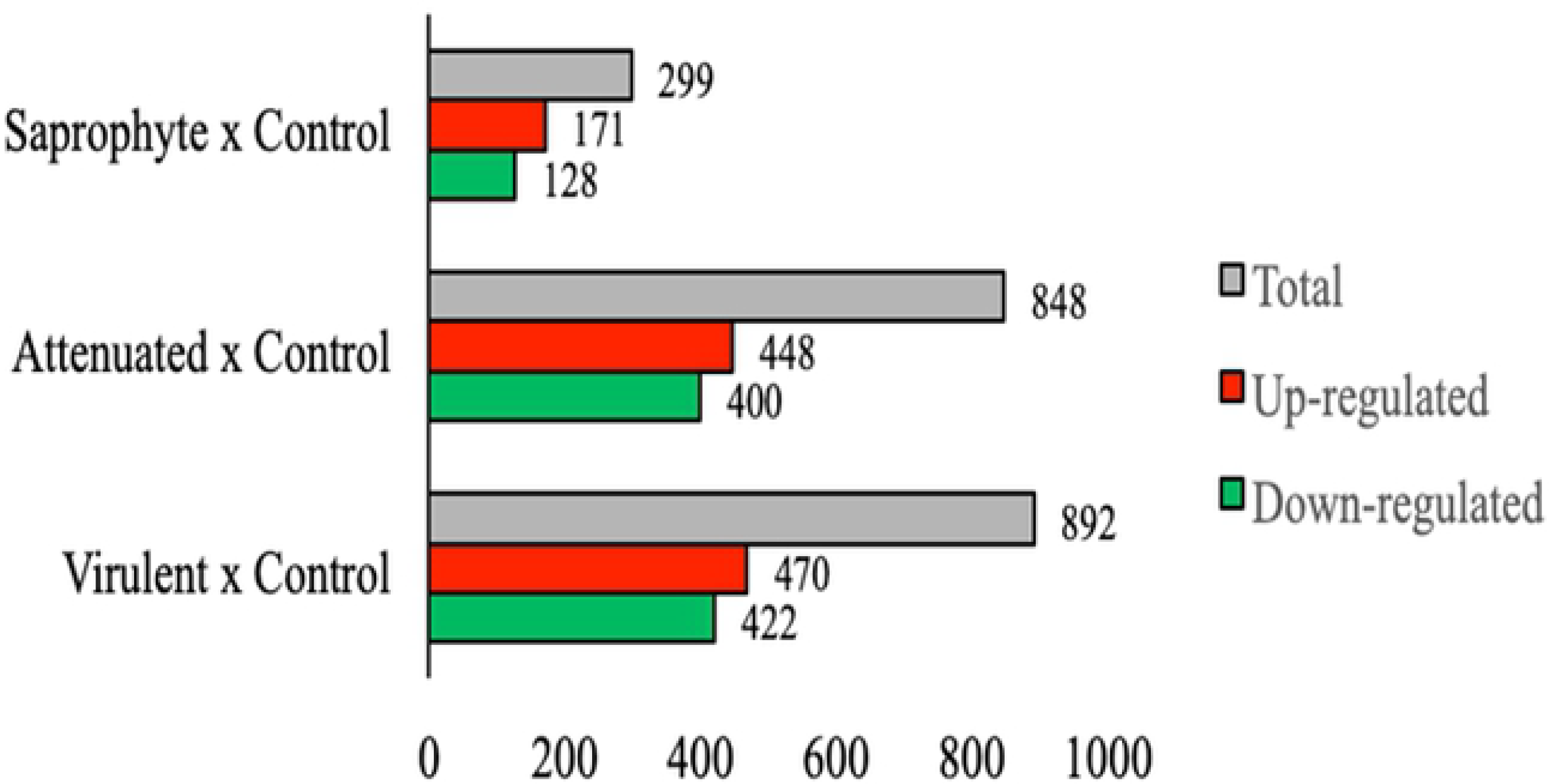
Differentially expressed genes after 6 hours of infection in murine macrophages J774A.1 with saprophytic, culture-attenuated and virulent strains of *Leptospira* spp. The colored bars show the up-regulated (red) or down-regulated (green) genes and grey a total of genes. (n = 3 / assay, FDR-adjusted p<0.05, fold change ± 2).

**Figure 2.**
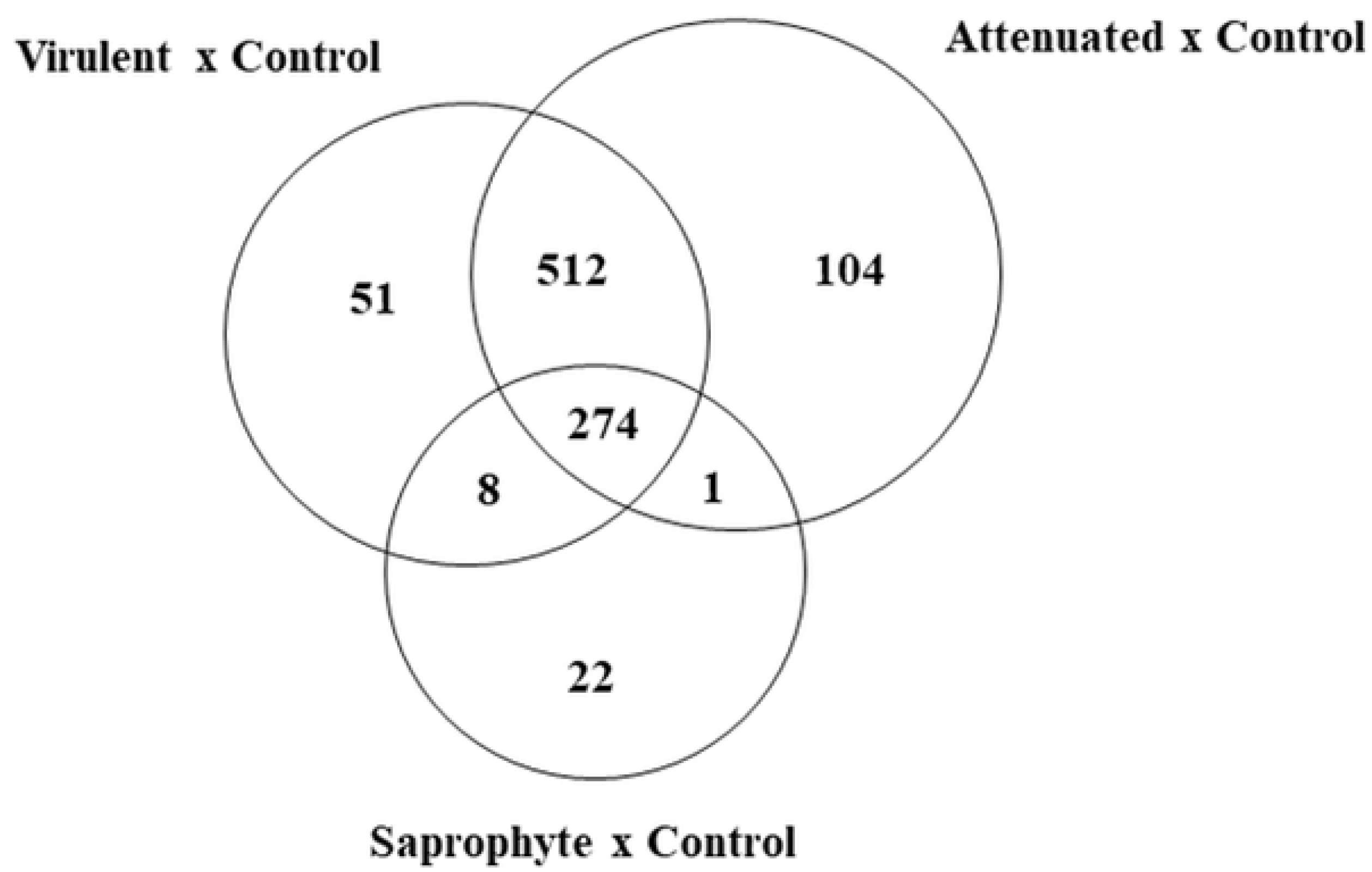
Venn diagram for differentially expressed genes of modulated macrophages at 6 hours of infection with different strains of *Leptospira* spp. Total number of canonical pathways (n=3/treatment; FDR<0.05, fold change ± 2) in the contrasts Infected (Saprophyte; Attenuated and Virulent) vs. Non-infected Control.

**Figure 3.**
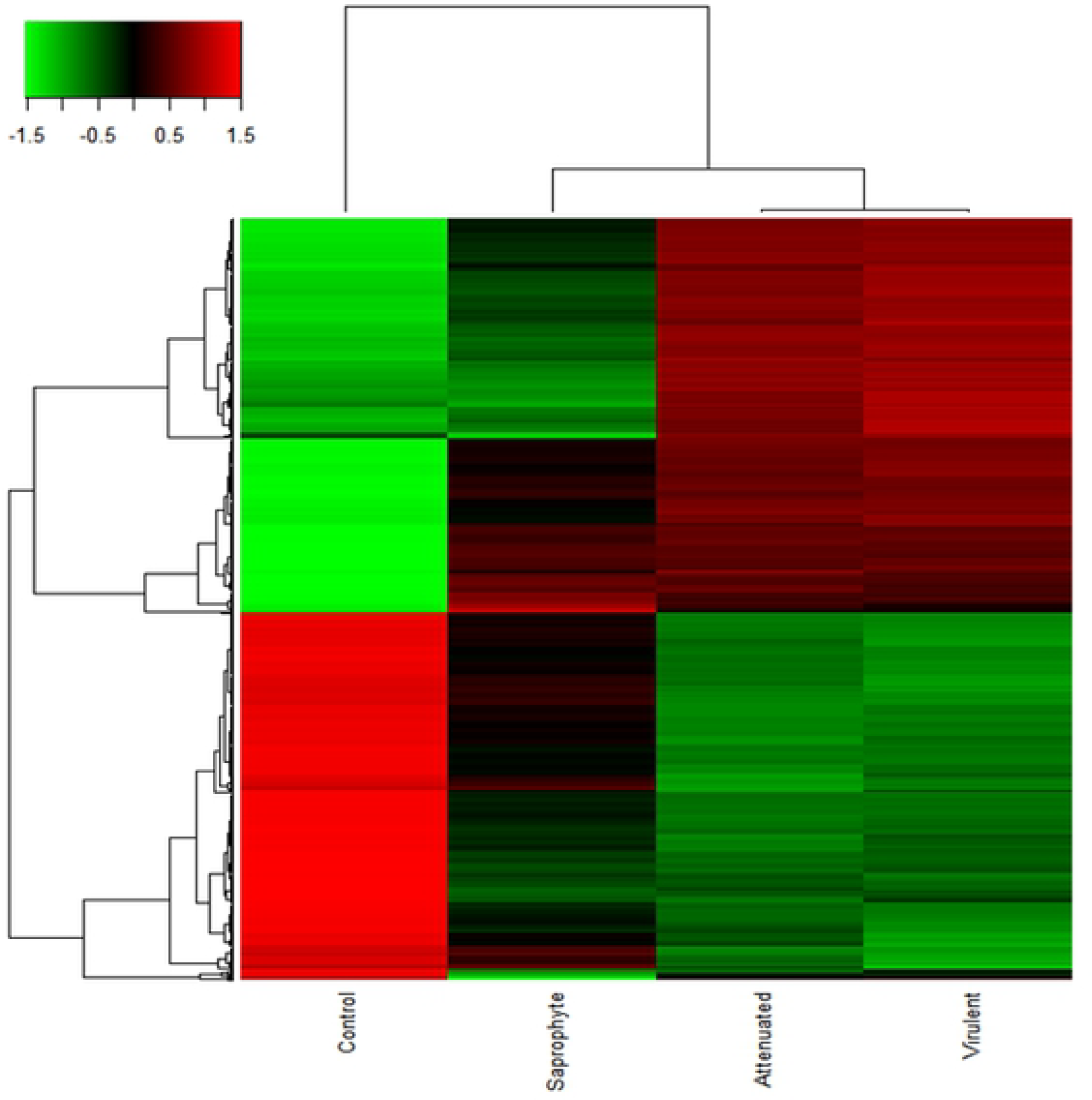
Heatmap of differentially expressed genes shows the average signal by macrophages at 6 hours of infection in different strains of *Leptospira* spp. The red color indicate increased expression, green color indicates the decreased expression as compared to control (*n* = 3/treatment; *p*-value < 0.01; FDR < 0.05; linear fold change ± 2).

In Table 1 we depict the top 9 DEGs in response to infection. These genes are present in several pathways and biological processes involved in acute inflammatory response.

**Table 1.** Top modulated transcripts in murine macrophages following 6h of infection with saprophyte, culture-attenuated and virulent strains of *Leptospira* spp.

| Regulation | Gene Symbol | FDR-adjusted p-value | FC (Sap vs. CT) | FC (Att vs. CT) | FC (Vir vs. CT) |
| --- | --- | --- | --- | --- | --- |
| **Up** | Il1b | 0,000037 | 202,85 | 253,12 | 259,03 |
|  | Il1a | 0,000004 | 42,58 | 120 | 127,28 |
|  | Saa3 | 0,000003 | 58,32 | 113,49 | 95,44 |
|  | Il6 | 0,000005 | 17,92 | 90,05 | 93,09 |
|  | Ccl5 | 0,000008 | 15,72 | 56,83 | 56,56 |
|  | Ptgs2 | 0,000011 | 38,21 | 52,72 | 56,33 |
|  | Nos2 | 0,000008 | 6,83 | 48,29 | 51,82 |
|  | Cxcl10 | 0,000035 | 8 | 45,54 | 51,25 |
|  | Ifit1 | 0,000217 | 6,16 | 33,6 | 37,61 |
| **Down** | Rasgrp3 | 0,000059 | -6,97 | -9,16 | -7,8 |
|  | Ighm | 0,000045 | -4,88 | -8,22 | -7,91 |
|  | Hal | 0,000022 | -4,52 | -6,78 | -7,01 |
|  | Cxcr4 | 0,000133 | -5,65 | -5,93 | -6,11 |
|  | Klhl24 | 0,000053 | -4,81 | -5,77 | -5,76 |
|  | Il18rap | 0,000074 | -3,7 | -5,55 | -5,31 |
|  | Nrcam | 0,000032 | -3,37 | -5,43 | -5,05 |
|  | Il1rl1 | 0,000028 | -3,24 | -5,35 | -4,53 |
|  | Ankrd44 | 0,000476 | -2,68 | -5,3 | -4,67 |

(FC=fold change; SAP= saprophyte; Att=attenuated; Vir=virulent)

### Analysis of signaling pathways

For functional enrichment of the differentially expressed genes obtained for each treatment, the Ingenuity Pathway Analysis (IPA) software was employed, Core Analysis was performed to identify relevant biological pathways to all 3 strains using the -log BH p-value > 1.3 (equivalent to a p-value <0.01).

### Specific pathways modulated by the virulent strain

Several pathways were identified as regulated by the virulent strain, however the Apoptosis pathway, ATM signaling and Cell Cycle: G2 / M DNA Damage Checkpoint Regulation, were exclusively expressed and affected by treatment with the virulent strain (Fig. 4). In the apoptosis pathway, the major up-regulated genes were FAS, IKBKE, NFKB1, NFKBIA, NFKBIB, NFKNID, NFKBIE, TNF, TNFRSF1B; downregulated transcripts were BCL2, CAPN2 and PARP1 (Fig. 5A). In the ATM signaling pathway, the upregulated transcript genes were CDKN1A, GADD45G, MDM2, NFKBIA and TLK2; downregulated transcripts were BRCA1, CBX5, CDK2, CHEK1, CHEK2, FANCD2, MDC1 and TOPBP1 (Fig. 5B). The upregulated genes of the Cell Cycle: G2 / M DNA Damage Checkpoint Regulation pathway were CDKN1A and MDM2; downregulated transcripts were BRCA1, CHEK1, CHEK2, PKMYT1 and WEE1 (Fig. 5C).

**Figure 4.**
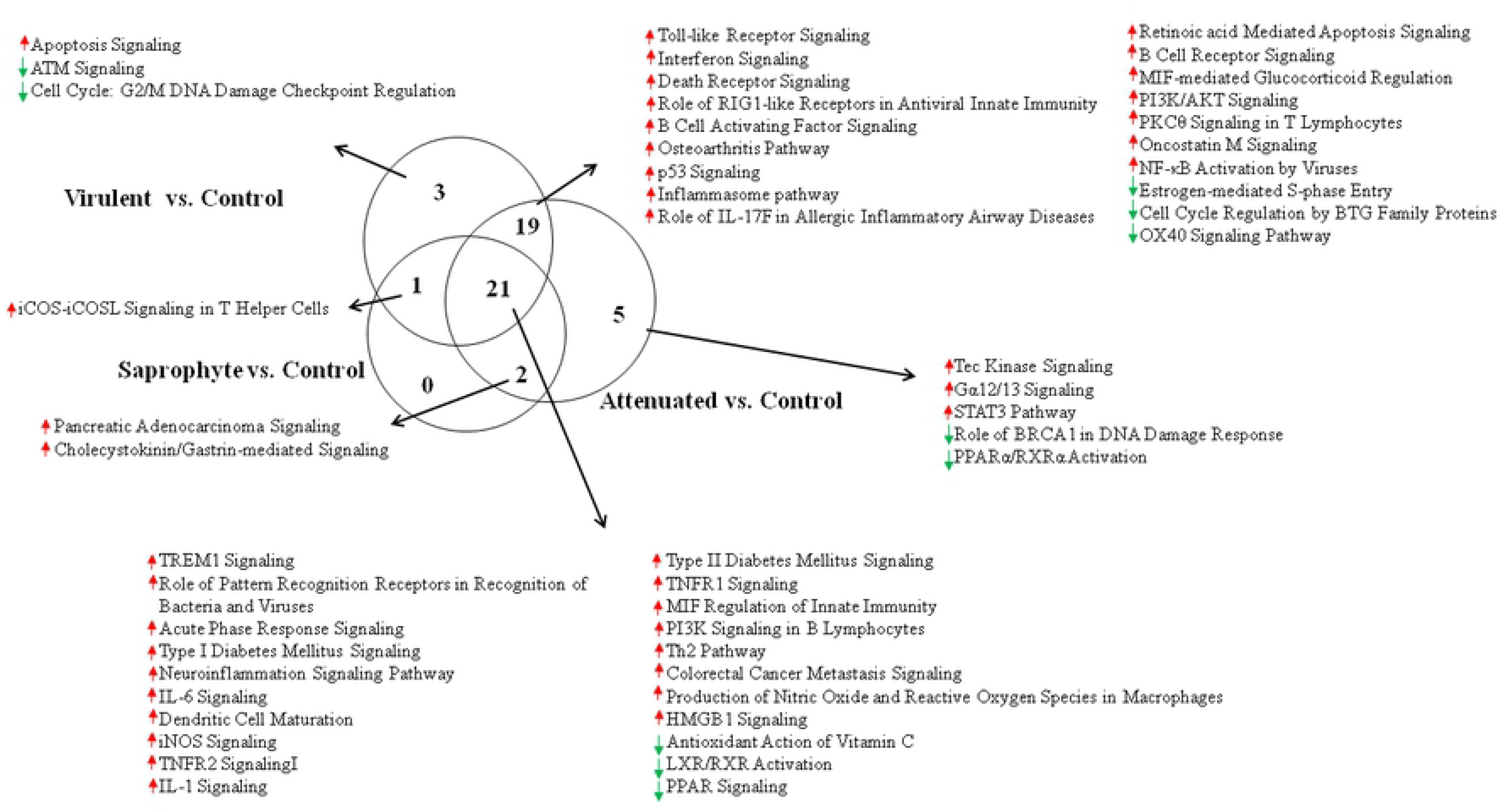
Venn diagram for pathways of modulated macrophages at 6 hours of infection with different strains of *Leptospira* spp. Total number of canonical pathways (n=3/treatment; FDR<0.05, fold change ± 2) in the contrasts Infected (Saprophyte; Attenuated and Virulent) vs. Non-infected Control.

**Figure 5.**
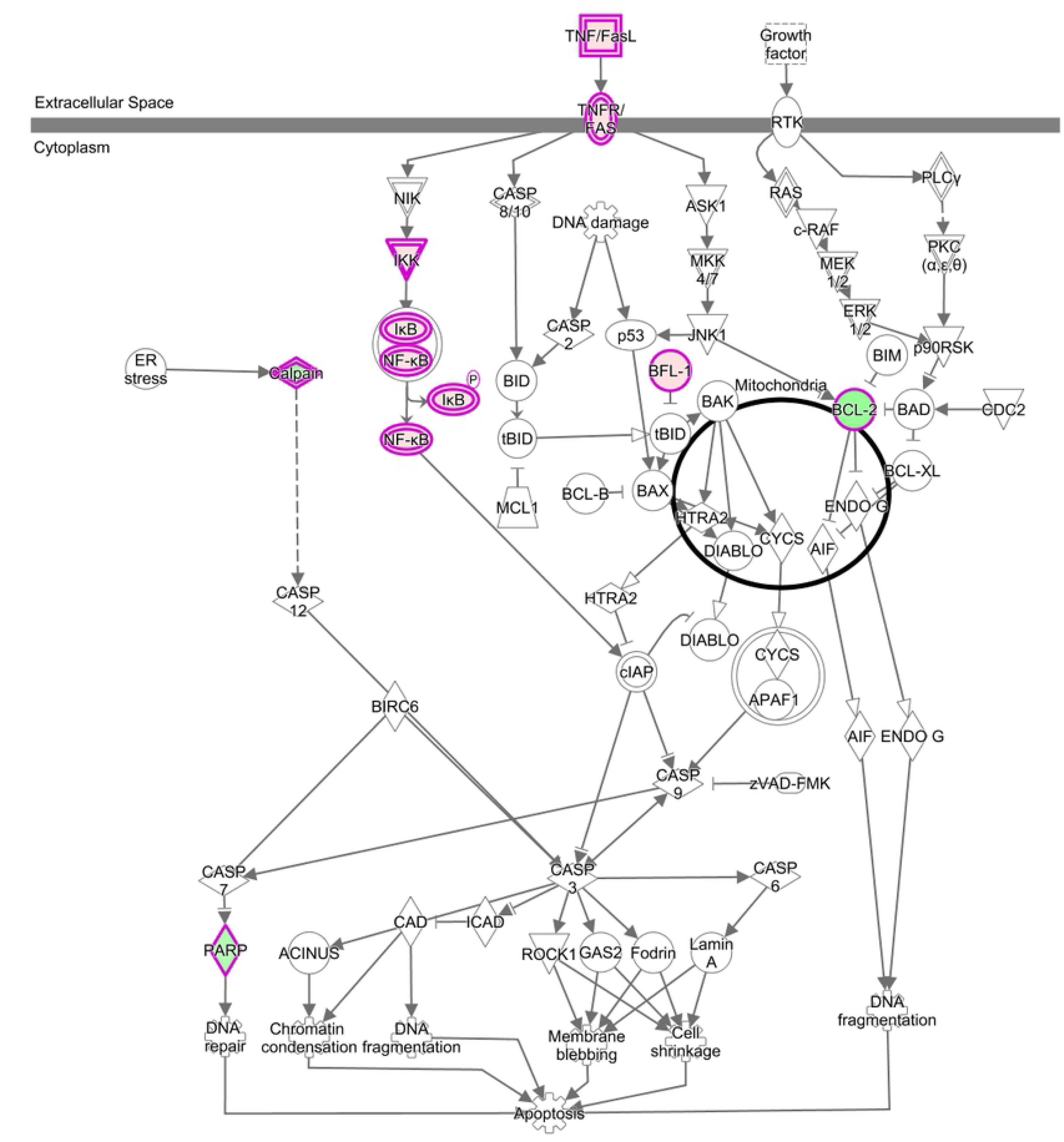
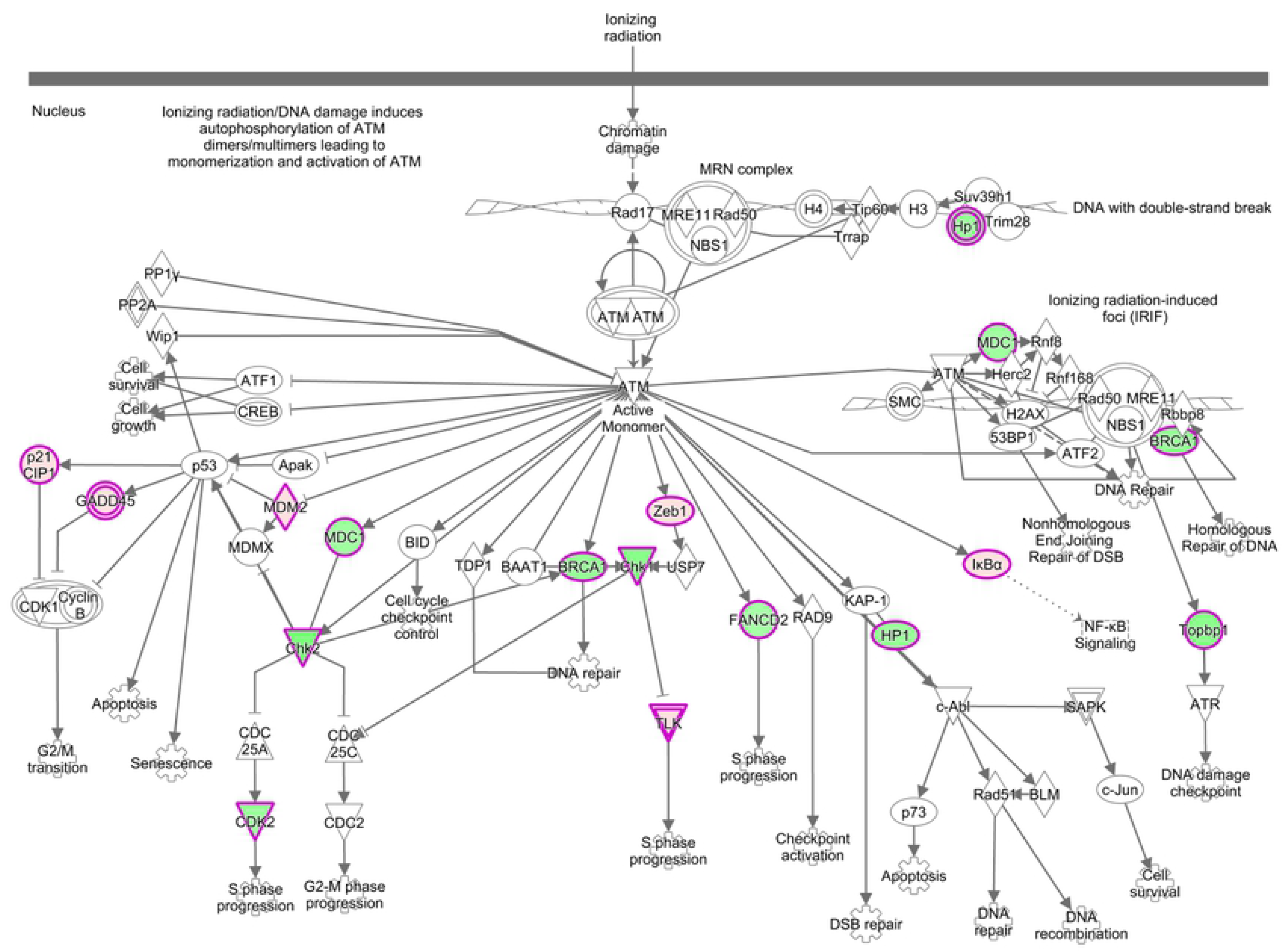
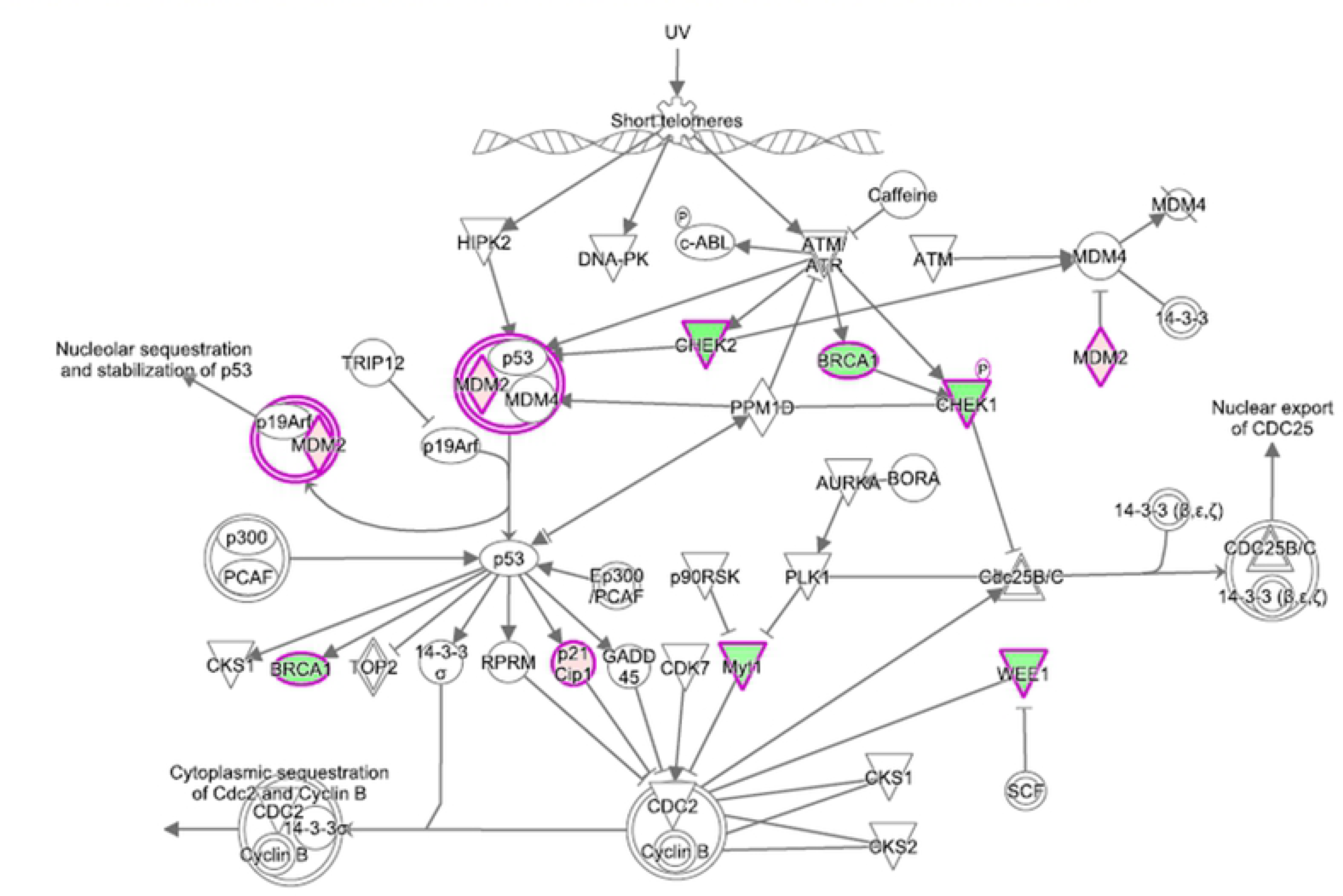
Canonical signaling pathway obtained by the IPA (Ingenuity Pathway Analysis program. Red and green indicate upregulated and down-regulated genes, respectively, compared to control group, and belongs to datasets of DEGs virulent vs. control assays. Color intensity corresponds to the degree of up or downregulation (fold-change). White represents the known genes of the pathway without identification in the transcriptomic analysis. **Panel A)** Canonical signaling pathway of Apoptosis of *in vitro* macrophages. **Panel B)** Canonical signaling of ATM of *in vitro* macrophages. **Panel C)** Canonical signaling pathway of Cell Cycle: G2 / M DNA Damage Checkpoint Regulation of *in vitro* macrophages.

### Validation of microarray data by qRT-PCR

Infection with 10^7^ of virulent and attenuated (*L. interrogans* serovar Copenhageni) and saprophytic (*L. biflexa* serovar Patoc), induced significant increase of TNF-α expression in murine macrophages (p <0.0001) compared to control. Regarding expression of IL-1β and NOS2, a similar expression profile was observed between Control and Saprophy, which differed from the profile found in the Attenuated and Virulent samples. The comparative analysis of the expressed values for IL-1β and NOS2 were statistically different between the assays, compared to the attenuated and virulent strains, differing when compared to the control groups and infection with the saprophytic strain. Differently from the observed TNF expression results, there was significant difference across all assays (Fig. 6A-C).

**Figure 6.**
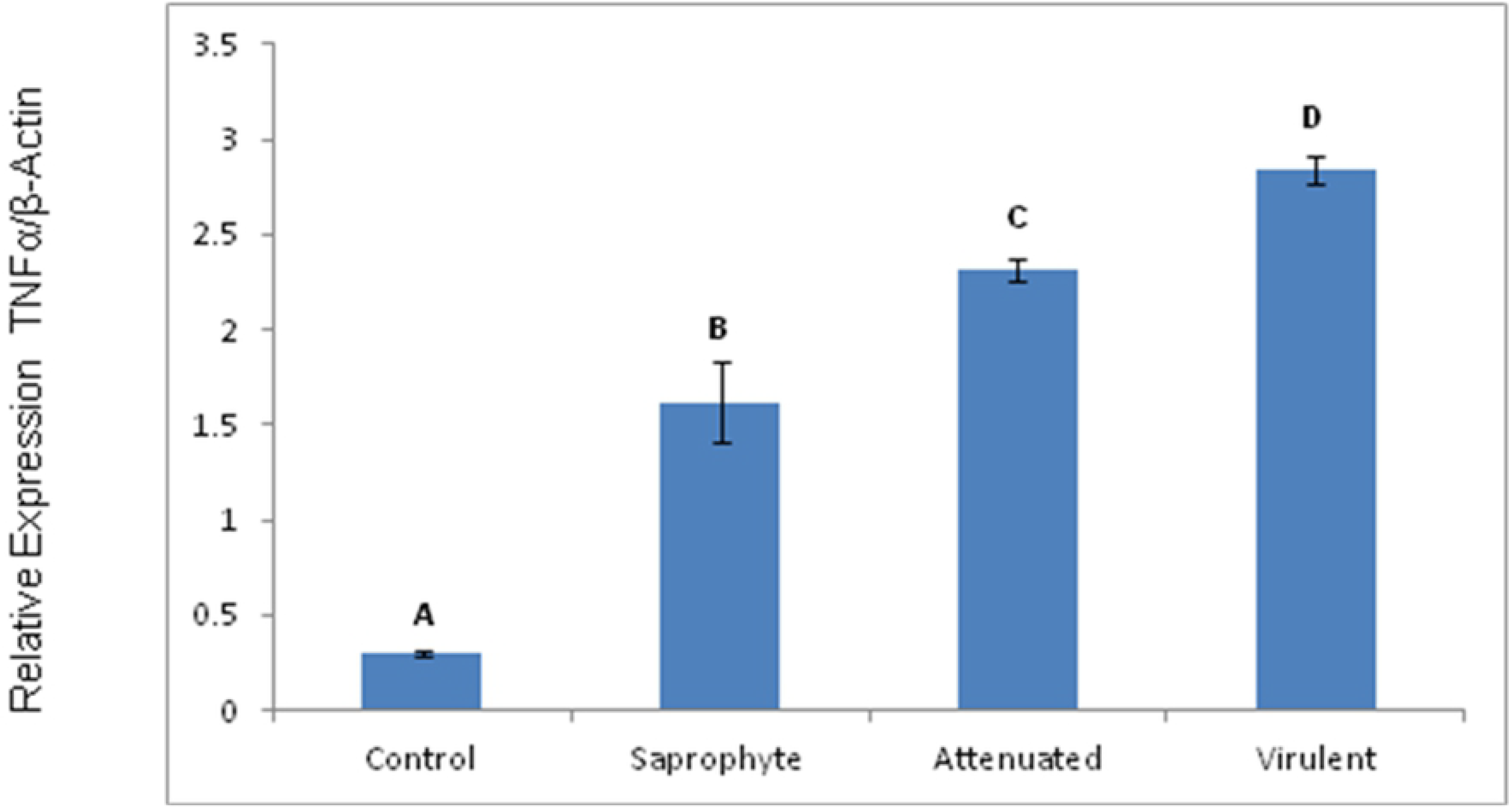
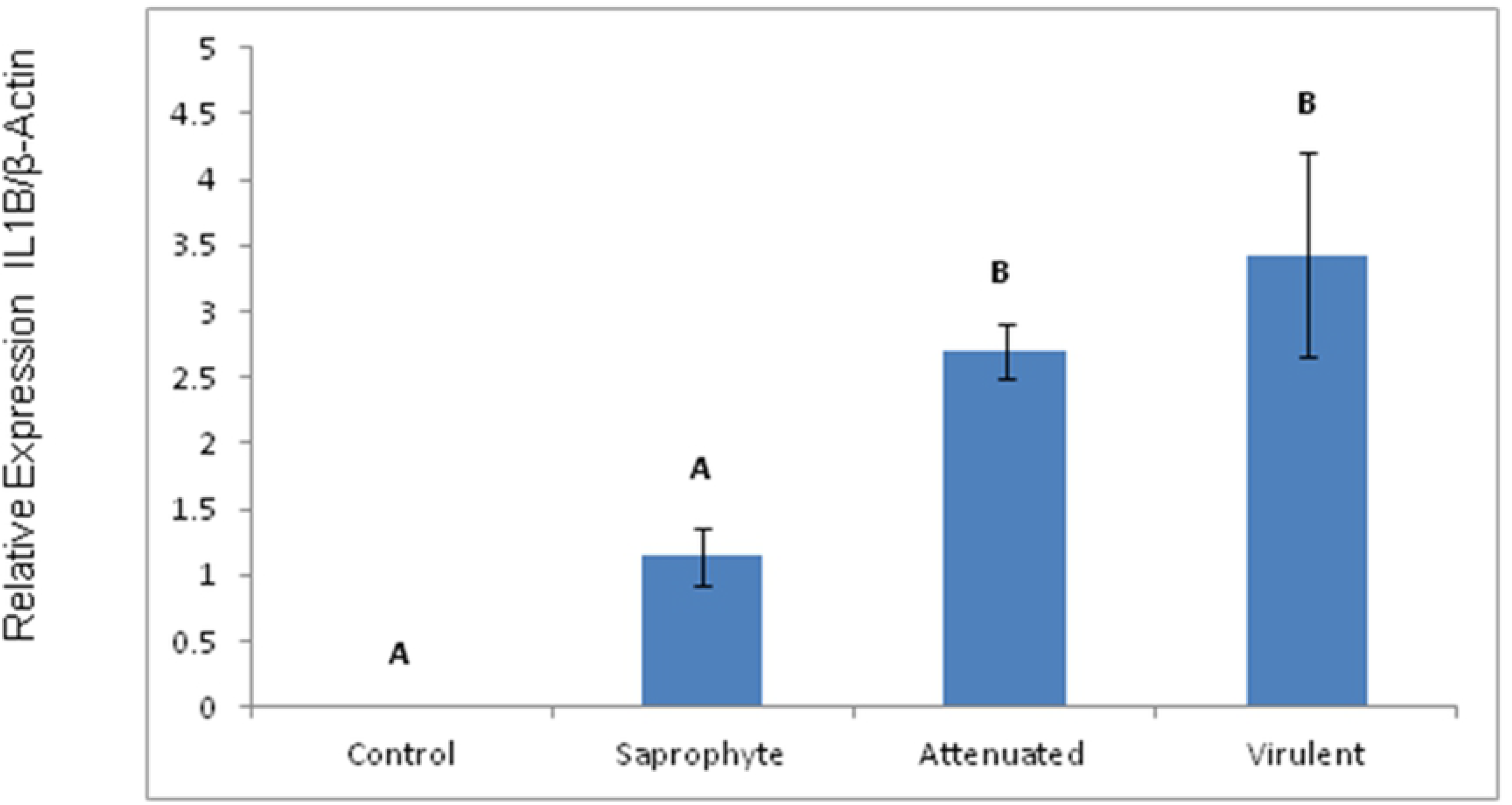
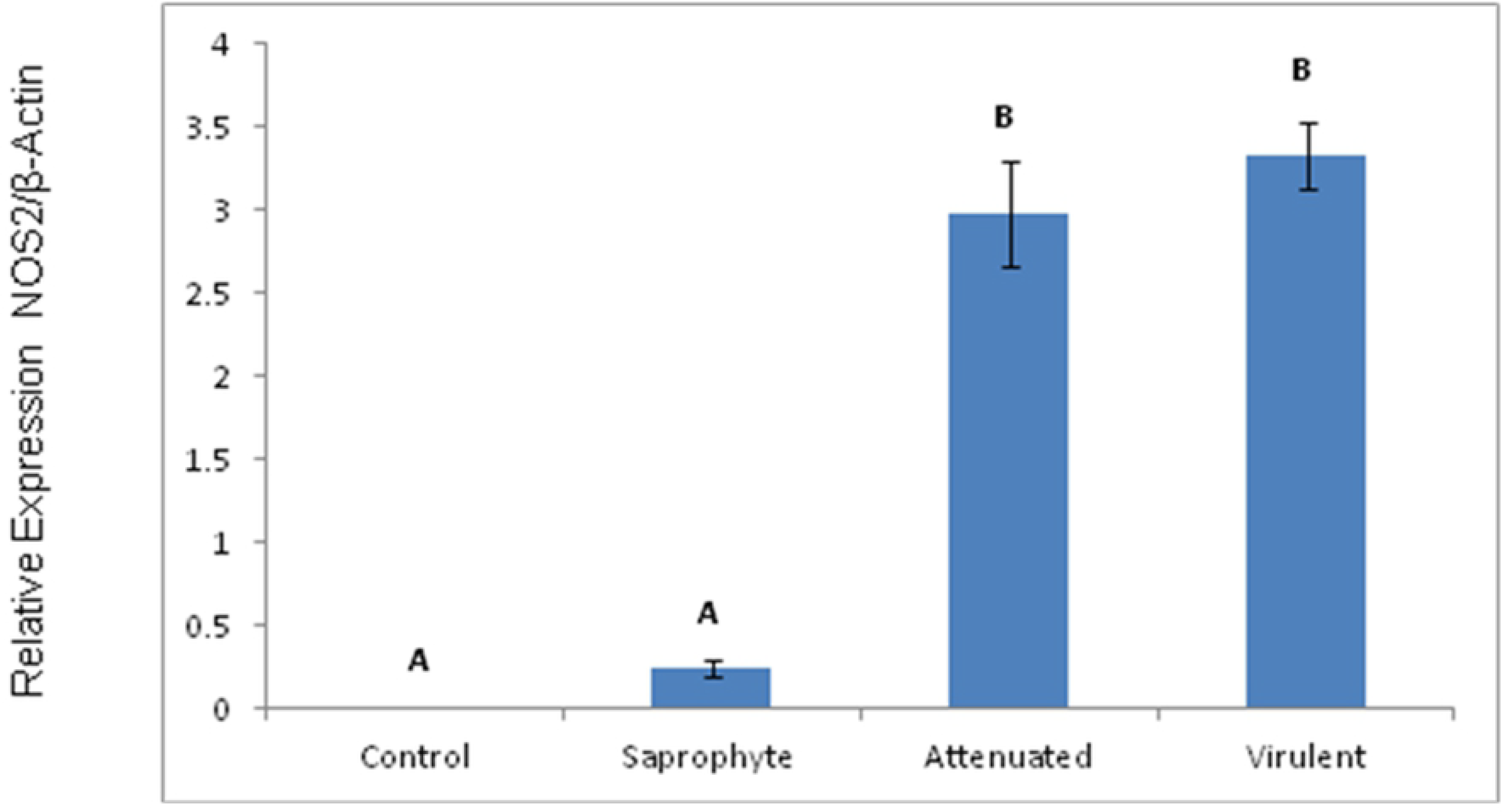
qRT-PCR of mRNA expression levels in infected macrophages with different strains of *Leptospira* compared to non-infected controls. **Panel A)** Relative expression of TNF-α in saprophyte, culture-attenuated and virulent compared to control. **Panel B)** Relative expression of IL-1β in saprophyte, culture-attenuated and virulent compared to control. **Panel C)** Relative expression of NOS2 in saprophyte, culture-attenuated and virulent compared to control (p <0.05). Different superscript letters differs significantly (p <0.05).

## Discussion

In this study, we took an *in vitro* approach to analyze the trancriptomic profiles of macrophages in response to saprophytic, culture-attenuated and virulent samples of *Leptospira* spp, to gain a better understanding of the disease’s molecular mechanisms and pathways.

TREM-1 signaling was the most significant pathway modulated by all threes strains. TREMs are a family of recently discovered receptors of the immunoglobulin superfamily, expressed on various cells of the myeloid lineage, which play important roles in innate immune responses, such as activating inflammatory responses and contributing to septic shock response in microbial-mediated infections [30–31]. Targeted activation of TREM-1 in our study appears to be a first line inflammatory response to the genus, regardless of virulence.

Other canonical pathways related to the innate immune system were a common response to all strains, the acute phase response signaling pathway, iNOS, IL-6, IL-1, TNFR1/TNFR2, MIF-regulation of innate immunity and HMGB-1 signaling. Further, all three strains are proposed to negatively regulate the antioxidant action of vitamin C pathway, suggesting that *Leptospira* spp. infection could contribute to oxidative stress associated production of reactive oxygen species (ROS). ROS-mediated intracellular oxidation is prevented by an antioxidant system, which includes low molecular weight antioxidants, such as vitamin C. This pathway is involved in cell process of survival, growth, proliferation and death [32].

In the culture-attenuated and virulent samples, Toll-like receptors, Interferon and inflammasome signaling pathways were significantly represented. Innate immune response is initiated by recognizing pathogens through pattern receptors as TLRs. Activation of these receptors is characterized by the massive production release of proinflammatory mediators, such as cytokines, chemokines and interferons [33]. Our microarray results to virulent strain identified high gene expression for TNF-α, IL-1β, IL-6 and iNOS. Similarly of Iskandar et al [34], verified that the presence of IL-6, IL-8 and IL-1β in serum from human patients with leptospirosis associated with severity of disease. In fact, Schulte et al. [35] concluded that increased TNF-α, IL-1β and IL-6 can activate the coagulation system in endotoxemic models. The high concentration of IL-6 is an indicator of septic shock and correlates with the severity of leptospirosis [36].

In regards to signaling pathways regulated by mRNAs modulated specifically following infection with virulent *L. interrogans*, Apoptosis signaling was positively regulated by infection whereas ATM signaling and Cell Cycle: G2/M DNA Damage Checkpoint Regulation, responsible for cell cycle, DNA repair and apoptosis, were negatively modulated. Following DNA damage, cells must detect breaks and transiently block the cell cycle progression allowing time for repair [37]. Jin *et al* 35 concluded that the pathogenic *Leptospira* caused apoptosis between 3-6 hours after infection. Our data corrobarates their finding that virulent *Leptospira* could modulated apoptosis, with just 6 hours of infection, through inhibition of pathways responsible for DNA repair and cell cycle control, as well as by inhibition the BCL-2 (anti-apoptotic gene) in turn leading to DNA damage and degradation. In fact, a previous study from our group has shown that BCL2 is a potentially down-regulated by mmu-mir-7667-3p following infection with *L. interrogans*, suggesting that cell survival could be compromissed after macrophages infection by the spirochete [38].

Leptospiral infection in macrophages induces a dependent p53/p21 cell cycle arrest [39]. We verified that the p53 target pathway signaling is regulated after virulent infection by modulation of mRNAs in murine monocyte-machrophages. Homotetrameric transcription factor p53 is reported to directly regulate 500 target genes, thereby controlling a broad range of cellular processes including cell cycle arrest, cell senescence, DNA repair, metabolic adaptation and cell death [40].

Further results from our study support the idea of cellular apoptosis Caspase-3 and 8 were elevated in all three infected macrophages, regardless of the pathogenicity notwithstanding, a more pronounced upregulation was induced by the with virulent and attenuated inoculum of *Leptospira*. Whether macrophage apoptosis induction by *Leptospira* is form of evasion mechanism or a host defense response to infection, preventing the spread of infection [jin 35] is still up for debate.

Cytokines represent a group of proteins that promote communication between cells, and their activation is through differentiation, receptor expression and cell-mediated immunity [41]. This suggests that the virulence factors, expressed or not during the process of infection of *in vitro* macrophages can guide cell response. In other words, virulent, culture-attenuated and non-pathogenic samples of *Leptospira* should be able to activate the murine macrophages, and the gene expression elicited as a result of infection, is dependent on strain virulence samples. Our results revealed a quantitative and qualitative association of gene expression with the virulence strains, with the virulent *L. interrogans* upregulating genes related to acute infection and cellular autophagy, unlike the culture-attenuated and saprophytic strains.

A comprehensive overview of gene expression patterns after infection by virulent, culture-attenuated and saprophytic *Leptospira* spp. strains revealed that inflammation and immune response, cytokine signaling, DNA repair, cell movement, death and cell survival were significantly activated following 6 hours of infection. Results demonstrated a group of genes is responsive to antigens present in the genus *Leptospira*, regardless of virulence, whereas species and virulence-specific gene expression was also elicited in the infected macrophages.

## Methods

### Leptospiral Strains

Samples of virulent strain *L. interrogans* sorovar Copenhageni (FIOCRUZ L1-130), attenuated strain *L. interrogans* sorovar Copenhageni M20 and saprophyte strain *L.biflexa* sorovar Patoc (FIOCRUZ - Patoc I) that we used in this study were donated by the Laboratory of Bacterial Zoonosis, Department of Preventive Veterinary Medicine and Animal Health of School of Veterinary Medicine and Animal Science, University of São Paulo (FMVZ/USP). All strains were incubated at 30ºC in Fletcher semi-solid culture medium.

### Macrophage Culture

Murine monocyte-macrophage cells (*Mus musculus* monocyte-macrophage cell line J774A.1), provided by the Paul Ehrlich cell bank (Rio de Janeiro, Brazil), was used as described [26]. Cells were maintained at 37º C, 5% CO2 in RPMI-1640 media (Sigma, USA) supplemented with 10% heat-inactivated fetal bovine serum (Gibco, USA), 100 ug/mL streptomycin (Sigma Chemical Co St.Louis, MO), 0.03% L-glutamine solution (Sigma) and 100 UI/mL of penicillin, in 6-well cell culture plates (3cm/well) until confluency [35].

### Infection of Macrophages

After the formation of confluent monolayer cells, they were washed three times in sterile phosphate buffer solution (pH 7.2) for removal of antibiotics and non-adherent cells. Bacteria were harvested by centrifugation and resuspended in RPMI-1640 media (Sigma). Cells were then infected (100:1 bacteria:cell) with *L. interrogans* L1-130 (virulent strain), *L. interrogans* M20 (culture-attenuated strain), *L. biflexa* Patoc I (saprophyte strain), as previously described [40]. Non-infected groups and non-infected macrophages were used as controls. All infected cells, in biological triplicates, were carried in fresh RPMI-1640, devoid of antibiotics, for 6h at 37º C, 5% CO_2_. Rate of infection did not differ between strains. At the end of the 6-hour period of infection, RNA extraction was immediately performed.

### RNA extraction and Quantification

Total RNA (n=3/experimental group) was extracted from macrophages with RNeasy Mini Kit (Qiagen, USA) according to manufacturer’s instructions. RNA samples were immediately stored at −80ºC. The quantification was performed using a NanoDrop (ND-2000 spectrophotometer, Thermo Scientific, Wilmington, DE, USA) and the samples quality was assessed using capillary electrophoresis (Bioanalyzer 2100 Agilent, Santa Clara, CA, USA). All samples used for microarray analysis had a RIN of 10.

### Transcriptome Array and Quality control

A WT PLUS Reagent Kit was used to prepare the RNA samples for whole transcriptome expression analysis with Mouse Genome 2.1 ST Arrays Strip Affymetrix (Santa Clara, CA, USA), according to the manufacturer’s protocols. Briefly, 100 ng of control RNA sample (Hela cells) was prepared to contain spiked in Poly-A RNA controls (lys, phe, thr and dap) absent in eukaryotic cells and mixed together with RNA samples to generate cDNA. After the amplification process, final cDNA was purified, quantified, fragmented and then labeled for hybridization to the strips, for 20h at 48ºC in the hybridization oven. Finally, strips were processed using the GeneAtlas Hybridization, Wash, Stain Kit for WT Array Strips (Affymetrix) and scanned using the GeneAtlas^®^ System (Affymetrix) generating the raw cell files. Raw intensity values in the cell files were background corrected, log2 transformed and then quantile normalized by the software Expression Console (Affymetrix) using the Robust Multi-array Average (RMA) algorithm.

### Identification of differentially expressed genes and functional enrichment

In order to identify differentially expressed genes, we utilized the software Transcriptome Analysis Console (Affymetrix), where statistical analysis was performed by one-way ANOVA (fold change ± 2, FDR corrected p<0.05). For the purpose of functional enrichment of the expression profiles obtained for each treatment, we used the Ingenuity Pathway Analysis (IPA) software (Qiagen).

### Validation of transcriptome results by qRT-PCR

For the validation of gene expression of selected genes in infected macrophages (saprophyte, culture-attenuated and virulent strains) and non-infected control macrophages, RNA samples were reverse transcribed (1µg of total RNA/sample) using the Moloney Murine Leukemia Virus (MML-V) enzyme (Life Technologies) and Oligo-dT Primers. All primers were designed to span at least one intron, to avoid repeat regions and similarities to other non-specific genomic regions. Mouse genome sequence, available on the University of California, Santa Cruz (UCSC) Genome Browser, was employed for primer design, using the Primer3 program [41]. PCR was performed using a Stratagene QPCR Systems Mx3005P (Agilent Technologies, Santa Clara, CA, USA) using the QuantiTect SYBR Green PCR kit (Qiagen). Expression levels were determined using standard curves for all genes at each individual run, and the expression of the candidate gene is presented as a ratio to an unregulated endogenous control (*β*-actin). The primers used for qPCR validation are listed in Table 2.

**Table 2.**
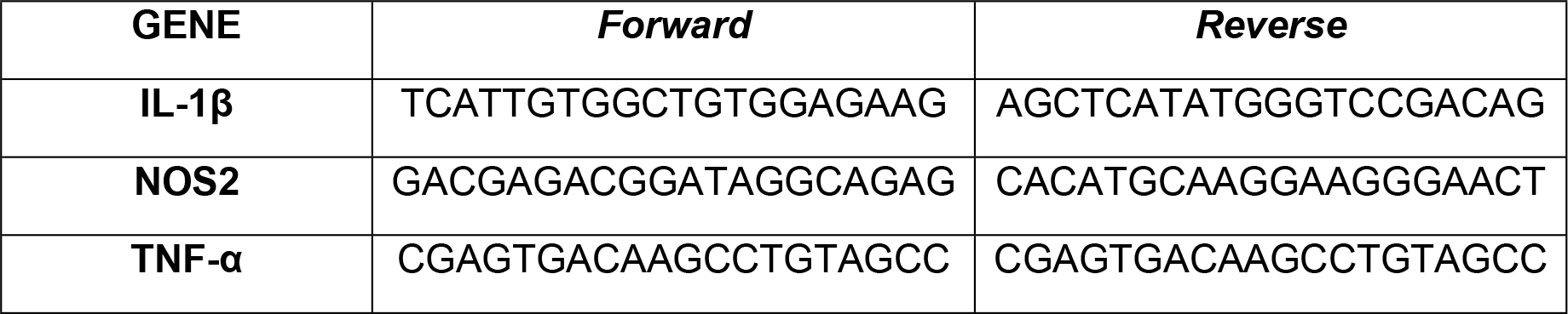
The primers used for qPCR validation.

### Statistical analysis

Differential expression of each gene was determined by one-way ANOVA with two criteria, a fold change of ±2 comparing all infected groups to the non-infected control and a Benjamini-Hochberg (BH) corrected p-value (FDR)<0.05. For pathways enrichment analysis on the Ingenuity Pathway Analysis (IPA) software (Qiagen), multiple testing was also (BH corrected (p<0.05). Real-time PCR data were analyzed using least-squares analysis of variance and the general linear model procedures of SAS (SAS Institute, Cary, NC, USA; p <0.01). Comparison of means was done using Duncan’s multiple range test.

### Ethics Statement

The present study was approved by the Research Ethics Committee of São Paulo State University (FMVA-UNESP), under the protocol number 2015-00895. No animal experimentation was performed in the experiments described herein.

## Acknowledgments

The authors would like to thanks Dr. Marcos Bryan Heinemmann by graciously providing the *Leptospira* spp., and to Cilene Vudovix Táparo for her valuable assistance during the experiments.

## Supporting information

**S1 Table. DEGs (gene symbol) modulated by macrophages at 6h of infection by different strains of *Leptospira* spp.**

